# Peripheral Neuropathy in the Adreno-myelo-neuropathy Mouse Model

**DOI:** 10.1101/2024.05.31.596769

**Authors:** Y Özgür Günes, C Le Stunff, JM Vallat, PF Bougnères

## Abstract

The Abcd1 knockout mouse mimics human adreno-myelo-neuropathy (AMN), thus contributes to a better understanding of disease mechanisms, which yet remain poorly identified. Only limited information is available about peripheral neuropathy (PN), although a notable component of AMN pathology besides myelopathy. To enrich our knowledge of PN, the current study reports the clinical, electromyographic and morphological aspects of peripheral neuropathy. We found that despite obvious electron microscopy anomalies in sciatic nerve axons, nerve conduction was nearly normal and did not seem to contribute significantly to the impaired motor performances of the Abcd1-/- mouse.

## Introduction

Adrenoleukodystrophy (X-ALD) is caused by mutations in the *ABCD1* gene located on the X chromosome which encodes a transporter that import very long chain fatty acids (VLCFA) into peroxisomes ^1^. The classical description of X-ALD is dichotomous and delineates two different clinical pictures. The more severe form, cerebral childhood adrenoleukodystrophy, (CCALD) affects the brain in late childhood as a rapidly progressive demyelination, leading to death before adulthood. The other neurological picture, adreno-myelo-neuropathy (AMN), begins between 20 and 50 years of age. AMN patients presents with spastic paraparesis and sensory ataxia, which aggravate progressively over the next decades, leading to severe spastic paraparesis. The majority of AMN neuropathology is attributed to a noninflammatory myelopathy due to a demyelinating axonopathy with axonal loss affecting some specific ascending and descending spinal cord tracts ^2^. Patients may also have PN, but, neuropathologic studies of peripheral nerves in AMN are scant and conflicting ^2-12^. Peripheral nerve lesions are variable and mild ^2^. Dorsal root ganglia (DRG), spinal nerve roots, sciatic, popliteal and ulnar nerves have been reported to be unremarkable in patients affected with manifestations ^2,5,6^, while sural and peroneal nerves have displayed a loss of large and small diameter myelinated fibers, endoneurial fibrosis, and thin myelin sheaths.

The Abcd1 knockout mouse closely mimics AMN and allows for investigating pathogenic mechanisms ^13^. Indeed, Abcd1-/- mice show no detectable clinical phenotype up to 15-18 months. Between 18 and 24 months of age, they progressively exhibit an abnormal neurological phenotype ^13^ characterized by impaired locomotor activity and coordination. These manifestations can be explained by myelopathy, since myelin and axonal anomalies are detectable in corticospinal tracts of mouse spinal cord ^13^. PN could also contribute to the impaired motricity and balance, since slower nerve conduction and myelin anomalies are observed in sciatic nerve ^13^. Our knowledge of PN, however, is based on only six Abcd1-/- mice that showed a large inter-individual variability ^13^. Preclinical gene therapy approaches of AMN have recently been attempted in Abcd1-/- mice with two different AAV9 vectors carrying the ABCD1 transgene. The first attempt targeted neurons that project to spinocerebellar or corticospinal axons respectively ^14,^ The other gene therapy vector used the MAG promoter to target oligodendrocytes and astrocytes in the white matter of the spinal cord^15^. None of these two studies targeted PN. Our belief is that PN should not be neglected when designing gene therapy strategy as it may contribute to the motor deficits of AMN patients. It is thus important to know more about the dysfunction and anomalies of the peripheral nervous system in the AMN mouse model.

## Results

Most of the studied 24-month old Abcd1-/- mice (N=18) showed impaired motor balance when tested with the crenelated bar (4.3 ± 1.1 vs 3.4 ± 0.8 in the wt, p<0.05) (Figure 1a). There was a large inter-individual variability across the mice, ranging from an almost intact (N=1, score 2) to a severely abnormal motricity (N=6, score ≥ 5), as previously reported at this age ^16^.

**Figure 1:**
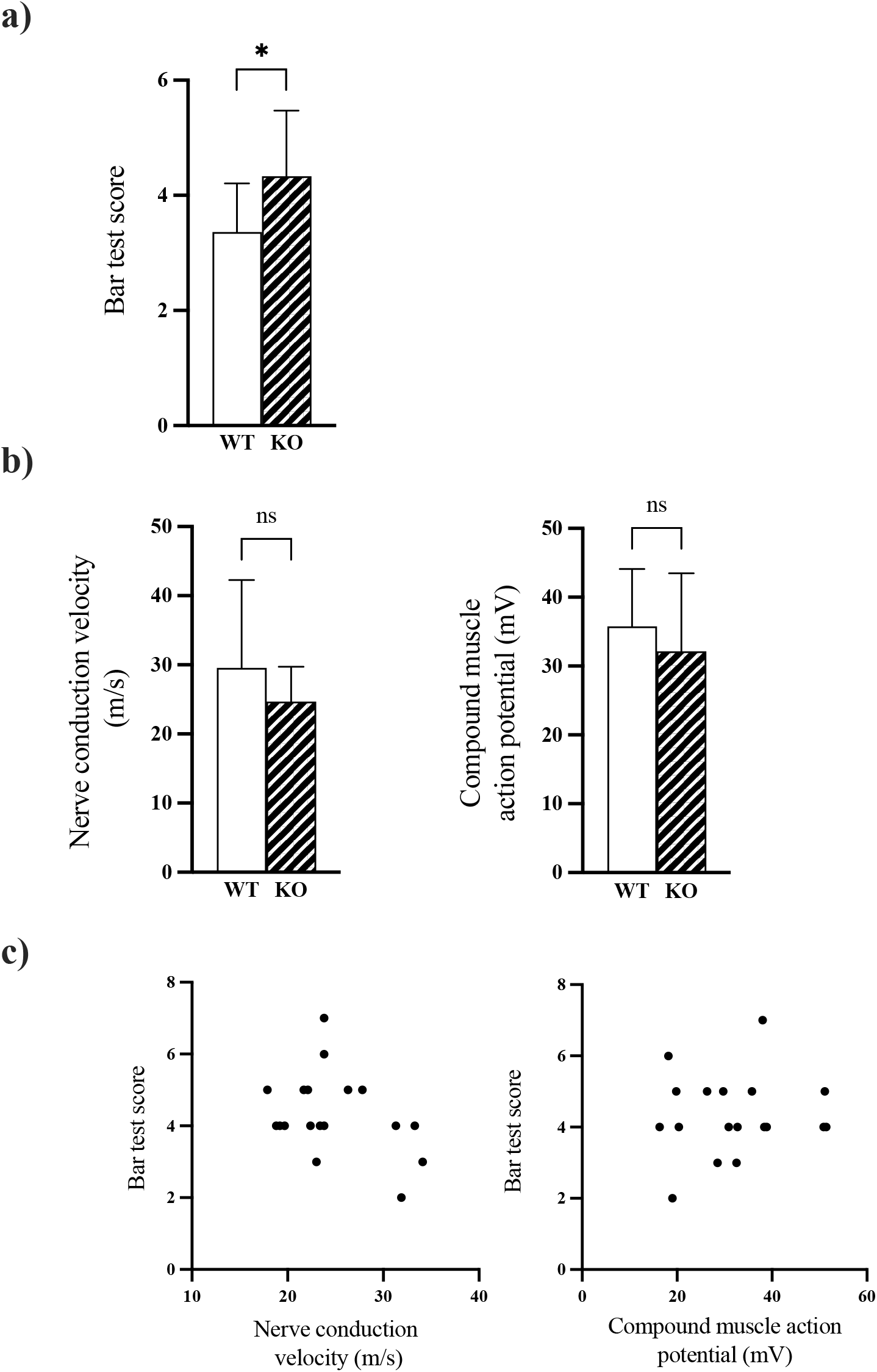
a) Bar test score is lower in Abcd1 knockout (N=18) compared with WT (N=8) mice p<0.05), b) Conduction velocity and compound muscle action potential (cMAP) in the same two groups of mice, (* p<0.05), c) Absence of correlation of bar test score versus NCV (r= 0.15, p=0.35) or CMAP (r= 0.097, p=0.70).

In contrast, the average conduction rate value in the sciatic nerve was only slightly decreased versus wt mice. Nerve conduction velocity averaged 24.7 ± 5.1 m/s ranging from 17.9 to 34.1 m/s compared to 29.6 ± 12.7 m/s in wt mice (NS) (Figure 1b). Compound muscle potential averaged 32.2 ± 11.3 mV ranging from 16.3 to 51.4 mV compared with 35.8 ± 8.4 mV in wt mice (NS) (Figure 1b). Given the inter-individual variability despite a strict experimental protocol and the limited size of mouse groups, the trend for a decrease of the two studied parameters did not reach statistical significance (P=0.34 for NCV and P=0.38 for CMAP). No correlation was found between NCV or CMAP and motor test (Figure 1c).

In the proximal sciatic nerve of Abcd1-/- mice, electron microscopy showed scant lesions of demyelination and limited axonal atrophy (Figure 2 a, b, c). Inflammation was not observed as previously reported ^2^. Schwann cells showed no expression of Aldp, while it was abundantly expressed in wt mice (Figure 3a, b).

**Figure 2.**
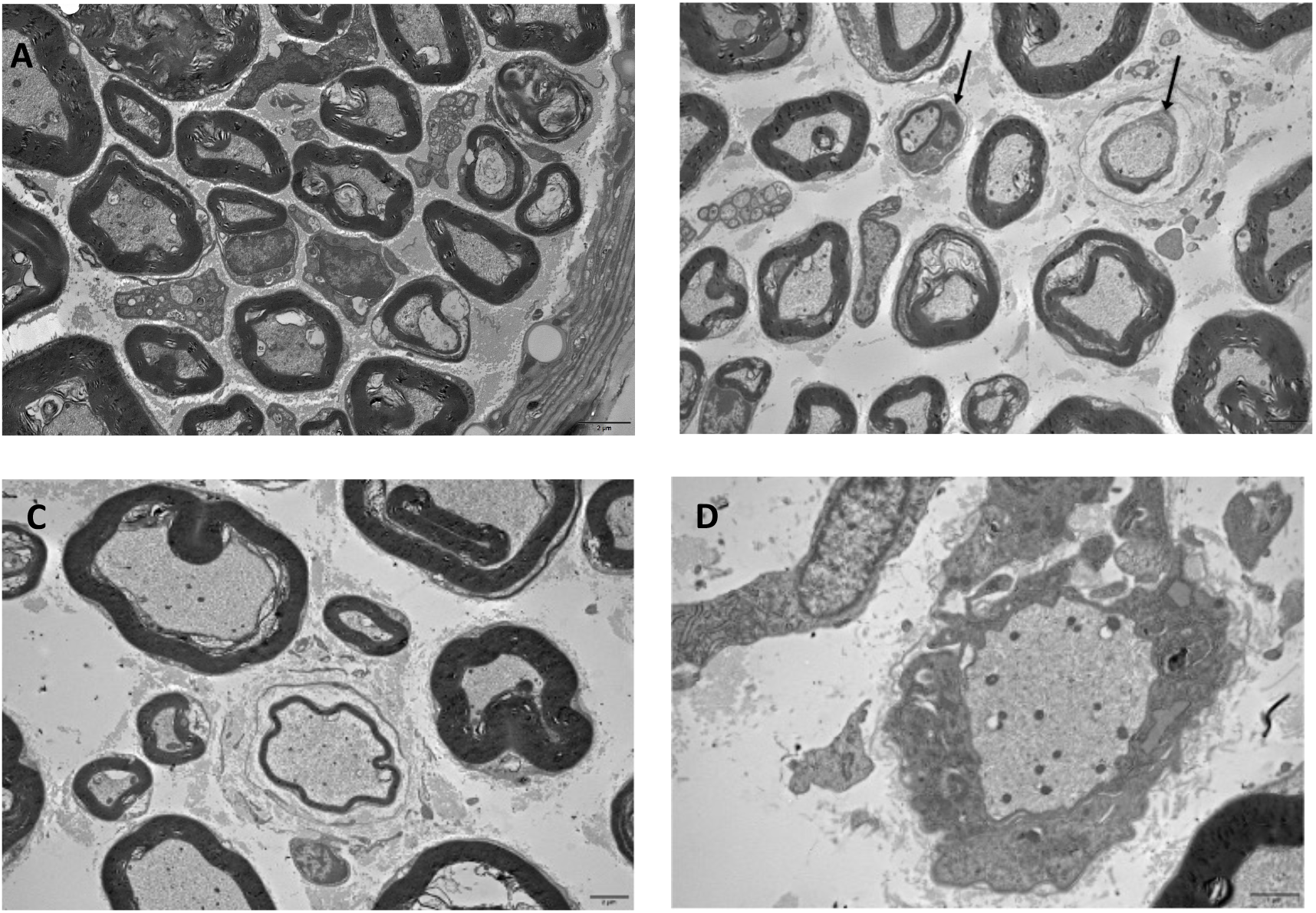
Electron microscopy images of sciatic nerve coronal sections in wt (A) and Abcd1 -/- mice (B-C-D). **A**. Normal density of myelinated fibers. No qualitative abnormality of these axons **B**. Two axons(arrows) have too thin myelin sheaths regarding their diameters, which indicates a demyelinating-remyelinating process. Scale bar 2µm. **C**. Another myelin abnormality surrounded by a concentric proliferation of Schwannian cytoplasm. Scale bar 2µm. **D**. Completely demyelinated axon. Scale bar 1µm.

**Figure 3.**
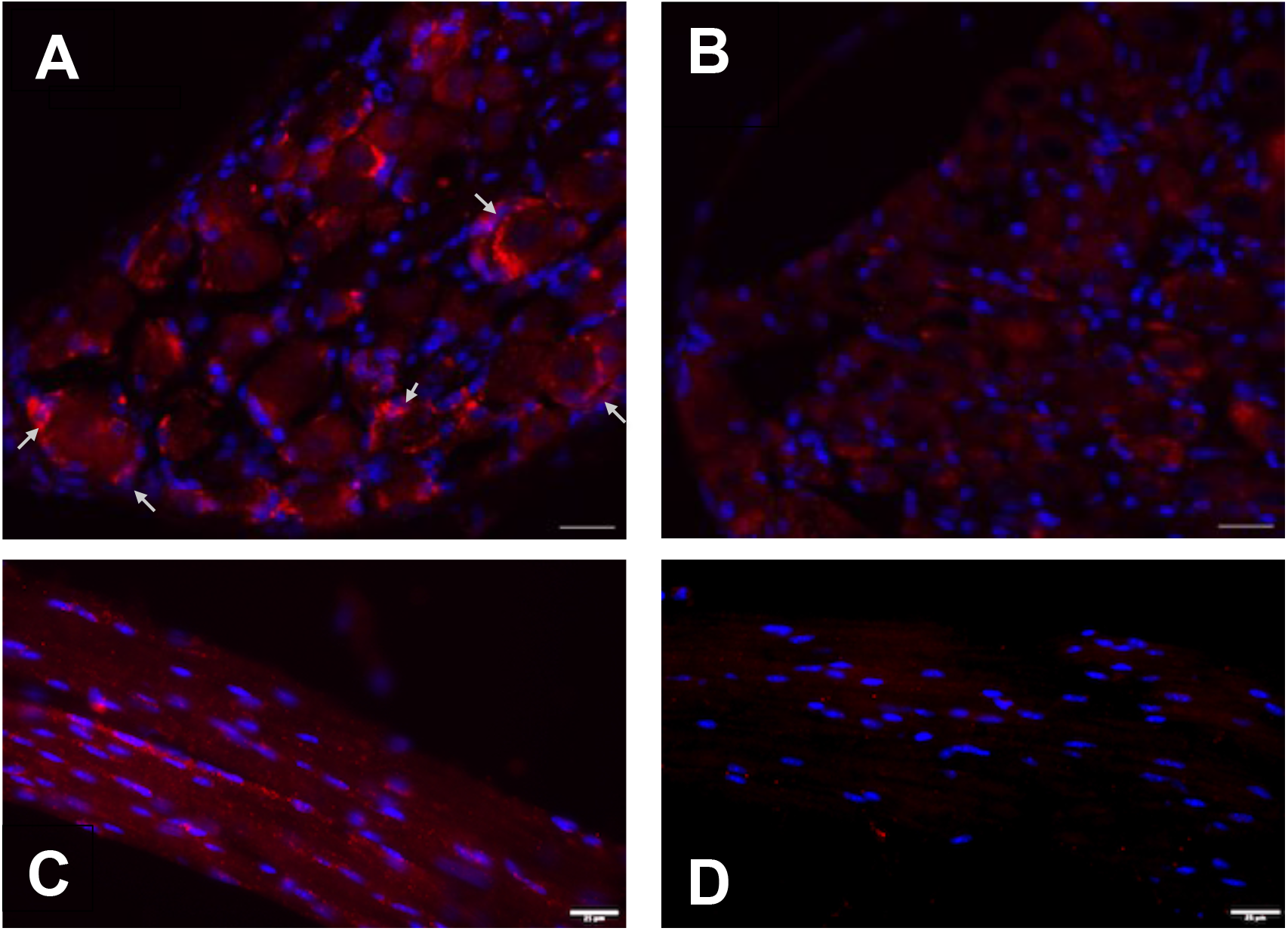
Mouse lumbar dorsal root ganglion. Satellite glial cells (arrows) surrounding cell bodies of large neurons express endogenous Aldp in wild type (A) not in Abcd1-/- mice (B). Dorsal root fibers expressing Aldp in wt (C), not in Abcd1-/- mice (D). Aldp (red); DAPI (blue).

In the DRG of Abcd1-/- mice, we found no expression of Aldp in neurons or satellite glial cells, while it was abundantly expressed in satellite cells of wt mice (Figure 3c).

## Discussion

A very small number of AMN patients ^2,3,5,6,8-12^ were reported to have microscopic examination of their peripheral nervous system. Some loss of myelinated fibers was observed in the lumbar plexus, and to a lesser degree in the brachial plexus. The femoral nerve showed similar changes. Electron microscopic of lumbar DRG revealed thinly myelinated fibers with some abnormalities of Schwann cells ^2^. Electromyographic studies revealed a mild impairment of nerve conduction in 16/23 AMN patients ^7^. For obvious ethical reasons, correlation between peripheral neuropathology and clinical manifestations could not be studied in patients.

The Abcd1-/- mouse model provided a synchronous description of functional and morphologic changes in the peripheral nervous system at a time of impaired motricity. The fact that we found a complete lack of correlation between nerve conduction parameters and motor test strongly supports that peripheral neuropathy has only a limited, if any, contribution to the impaired motricity. For example the mouse having the worst combination of EMG parameters (19.2 m/s and 20.4 mV-21,7 m/s) had a bar score of 4, similar to the mouse who had the best combination of EMG parameters (33.3 m/s and 38.4 mV). The mouse with the worst bar score at 7 had unremarkable EMG parameters at 23.8 m/s and 38 mV.

The morphological abnormalities in myelin structure and axons that we observed in the sciatic nerve add little to previous reports ^13^. Our microscopic analysis was only qualitative, but clearly showed that no relationship exists between morphological changes in sciatic nerve and EMG parameters or motor performances.

The studied Abcd1-/-mice showed a complete lack of Aldp expression in the Schwann cells of the proximal sciatic nerve, while wt mice expressed abundant Aldp. The lack of Aldp expression was also observed in DRG neurons and satellite cells, while many satellite glial cells expressed Aldp, as reported ^17,18^.

In conclusion, the functions of the peripheral nervous system seems only mildly affected in Abcd1-/- mice. Our study also opens the door to testing the effect of gene therapy approaches in this model, notably with regard to the restoration of Aldp expression in Schwann cells and cells of the DRGs.

## Materials and Methods

### Animals

C57BL/6 Abcd1-/- and wild-type (WT) C57Bl/6 mice were backcrossed onto a pure C57/B6 background over three generations, then bred from homozygous founders, and occasionally genotyped. They were kept in the animal housing facility of the MIRCen Institute, had ad libitum access to water and standard food, and were kept on a 12-h light/12-h dark cycle. Experiments were approved by the Institutional Animal Care and Use Committee at MIRCen CEA and the Ministry of Research (APAFIS Nos. 02015040710445631 and 31476-2021051016324194 v1).

### Behavioral tests

Tests were conducted at 24 months of age. Animals were transferred to experimental room 30 min before each experiment. Mice were evaluated by behavioral testing by an examiner blind to the treatment condition. Videos of the mice were recorded during the crenelated bar cross tests, then scored by three investigators (CLS, YOG, and PB) blind to mouse condition. Technical details of the crenelated bar cross test are described in our previous study^15^.

The bar is just wide enough for mice to stand on with their hind feet hanging over the edge such that any slight lateral misstep will result in a slip. A score was defined by several qualitative and quantitative parameters more or less characteristic of each mouse. A normal mouse runs rapidly along the bar in a single stroke, without stopping, progressing in short leaps or by alternately advancing its front and rear erect legs, keeping its tail straight in line with a horizontal back, without slipping significantly except by excessive speed. In contrast, a severely diseased mouse shows a number of important hind paw fails (when one or both hind paws misses the merlon and slip in the crenelated parts of the beam), drags its hindquarters while using its front paws to move forward on the bar, maintain a hunchback attitude, use its curved, downward-pointing tail for balance, hesitate or stop for a moment and cover the bar in a prolonged time. The defect observed mainly concerns the motor skills of the hind legs, while the front legs seem to function normally.

The score of the bar test was assessed on a scale of 1–8, with 8 being the maximum aggregate score for the most diseased mice and 4 being a threshold for quoting a serious motor deficit. Videos of mice provided as supplemental data were taken by an observer who was not blinded to treatment. There was perfect agreement between the score values initially given by the three blinded investigators in 87% of videos, in 7% of cases the discrepancy was ≤ 0.5, in 4% of cases it reached 1, and in 2% of cases 2. In the event of even the slightest discrepancy, the video was re-examined at a later time, and if the discrepancy persisted (5% of cases), the values were averaged for the score finally assigned to the mouse

### Electromyographical studies

Electrophysiological recordings were performed on the sciatic nerve, according to ^19^.

Briefly, the animals were anesthetized with isoflurane. Measurements took about 30 min/animal in total. A constant body temperature of 37 °C was carefully maintained by a heating plate and continuously measured by a rectal thermo probe. Sufficient anesthesia was confirmed by testing simple reflexes such as movement reflexes and testing of sensitivity for low-grade pain. Ring electrodes were placed using contact gel for optimal conductivity / transfer resistance. The sensing electrode was placed at the position where the gastrocnemius muscle has its maximum diameter. Proximal and distal nerve stimulations were performed, registering the transmitted potentials with sensing electrode. After data processing, consistent values for the nerve conduction velocity (NCV) and the compound motor action potential (CMAP), the key parameters for quantification of gross peripheral nerve functioning, were achieved.

### Tissue preparation

Mice aged 24 months received a lethal dose of pentobarbital (Exagon, Axience, 400 mg/kg) by intraperitoneal injection. Animals then underwent intracardiac perfusion of 1X PBS followed by 4% paraformaldehyde in 0.01 M PBS for histological analysis or with 4% paraformaldehyde/2.5% glutaraldehyde for electron microscopy. Tissues were collected, postfixed overnight in 4% paraformaldehyde at +4°C, and cryoprotected in incrementing PBS-sucrose gradients before histological processing. Ten-micrometer-thick sections were collected by Leica CM3050 SC cryostat.

### Immunofluorescence staining

Following antibodies and reagents were used for immunofluorescence staining: anti-ABCD1/ALDP (Abcam, ab197013), biotin anti-rabbit (BA-1000; Vector Laboratories), Streptavidin-Cy5 (434316; Invitrogen), DAPI solution (564907; BD Biosciences), mounting medium (Vectashield; H1000; Vector Laboratories), (NGS) (S1000; Vector Laboratories), (BSA) (A9647; Sigma), Triton X-100 (Sigma).

